# Regulation of airway fumarate by host and pathogen promotes S. *aureus* pneumonia

**DOI:** 10.1101/2025.04.10.647998

**Authors:** Ying-Tsun Chen, Zihua Liu, Dario Fucich, Stefano G. Giulieri, Zhe Liu, Ridhima Wadhwa, Gustavo Rios, Henning Henschel, Nupur Tyagi, Françios A. B. Olivier, Ian R. Monk, Shivang S. Shah, Shwetha H. Sridhar, Marija Drikic, Colleen Bianco, Gaurav K. Lohia, Ayesha Z. Beg, Paul Planet, Ian A. Lewis, Robert Sebra, Ana Traven, Abderrahman Hachani, Timothy P. Stinear, Benjamin P. Howden, Jeffrey M. Boyd, Sebastian A. Riquelme, Chu Wang, Alice Prince, Tania Wong Fok Lung

**Affiliations:** Department of Pediatrics, Columbia University, New York, NY 10032, USA; Peking-Tsinghua Center for Life Sciences, Academy for Advanced Interdisciplinary Studies, Peking University, Beijing 100871, China; Department of Microbiology and Immunology, The University of Melbourne at the Peter Doherty Institute for Infection and Immunity, Melbourne, VIC 3000, Australia; Synthetic and Functional Biomolecules Center, Beijing National Laboratory for Molecular Sciences, Key Laboratory of Bioorganic Chemistry and Molecular Engineering of Ministry of Education, College of Chemistry and Molecular Engineering, Peking University, Beijing, 100871, China; Department of Microbiology, Biochemistry and Molecular Genetics, Rutgers New Jersey Medical School, Newark, NJ 07103, USA; Center for Immunity and Inflammation, Rutgers New Jersey Medical School, Newark, NJ 07101, USA; Department of Biochemistry and Microbiology, Rutgers University, New Brunswick, NJ 08901, USA; Department of Medicinal Chemistry, Uppsala University, Uppsala, Sweden; Infection Program and the Department of Biochemistry and Molecular Biology, Biomedicine Discovery Institute, Monash University, Clayton, VIC 3800, Australia; Department of Genetics and Genomic Sciences, Mt. Sinai Icahn School of Medicine, New York, NY 10029, USA; Department of Biological Sciences, University of Calgary, Calgary, T2N 1N4, Canada; Division of Infectious Disease, Department of Pediatrics, The Children’s Hospital of Philadelphia, Philadelphia, PA 19104, USA; Department of Pediatrics and Microbiology, Perelman School of Medicine, University of Pennsylvania, Philadelphia, PA 19104, USA; Panacent Bio, New York, NY 10014; Microbiological Diagnostic Unit Public Health Laboratory, The University of Melbourne at the Peter Doherty Institute for Infection and Immunity, Melbourne, VIC 3000, Australia

## Abstract

*Staphylococcus aureus* is a leading cause of healthcare-associated pneumonia, contributing significantly to morbidity and mortality worldwide. As a ubiquitous colonizer of the upper respiratory tract, *S. aureus* must undergo substantial metabolic adaptation to achieve persistent infection in the distinctive microenvironment of the lung. We observed that *fumC*, which encodes the enzyme that converts fumarate to malate, is highly conserved with low mutation rates in *S. aureus* isolates from chronic lung infections. Fumarate, a pro-inflammatory metabolite produced by macrophages during infection, is regulated by the host fumarate hydratase (FH) to limit inflammation. Here, we demonstrate that fumarate, which accumulates in the chronically infected lung, is detrimental to *S. aureus*, blocking primary metabolic pathways such as glycolysis and oxidative phosphorylation (OXPHOS). This creates a metabolic bottleneck that drives staphylococcal FH (FumC) activity for airway adaptation. FumC not only degrades fumarate but also directs its utilization into critical pathways including the tricarboxylic acid (TCA) cycle, gluconeogenesis and hexosamine synthesis to maintain metabolic fitness and form a protective biofilm. Itaconate, another abundant immunometabolite in the infected airway enhances FumC activity, in synergy with fumarate. In a mouse model of pneumonia, a Δ*fumC* mutant displays significant attenuation compared to its parent and complemented strains, particularly in fumarate- and itaconate-replete conditions. Our findings underscore the pivotal role of immunometabolites in promoting *S. aureus* pulmonary adaptation.

## Introduction

*Staphylococcus aureus* frequently causes healthcare-associated pneumonia (1). Antibiotic susceptible as well as resistant strains pose a significant risk for prolonged airway infection and associated morbidity (2). This risk is heightened in patient populations with underlying pulmonary pathology, such as chronic obstructive pulmonary disease (COPD), cystic fibrosis (CF), and following viral infection (3, 4). *S. aureus* is an especially formidable pathogen due to the expression of many surface proteins that thwart innate immune clearance (5). While toxin-producing epidemic isolates can cause a fulminant necrotizing pneumonia (6), typically the bacteria adapt to the airway milieu to cause a more indolent but intractable infection. These less immunostimulatory strains are better tolerated by the host as they fail to induce as robust an immune response that is detrimental to the host and pathogen alike. Of note, vaccine strategies that target classic virulence factors have failed (7), indicating the need for alternative approaches to control staphylococcal infection.

Among the common respiratory pathogens, *S. aureus* is notable for its substantial metabolic flexibility (8). In contrast to more fastidious pathogens, *S. aureus* readily adapts to local carbon sources to optimize its metabolism. The infected airway provides a unique microenvironment with a distinctive set of metabolites and molecules produced by both the host and pathogen including reactive oxygen species, and DNA from lysed or damaged immune and stromal cells. By studying patterns of gene expression in clinical isolates of *S. aureus* from patients with persistent pneumonia, we observed staphylococcal adaptation to airway metabolites that accumulate during ongoing infection (9). In a survey of methicillin resistant *S. aureus* (MRSA) strains isolated from chronic pneumonia, we noted over 100- to 1000-fold increases in the expression of *fumC*, which encodes the bacterial fumarate hydratase (FH) that converts fumarate to malate (10). Fumarate, a potent proinflammatory metabolite that induces the production of TNF and IFNβ in stimulated macrophages, is regulated by the host FH to limit immunopathology (11). While the immunoregulatory function of host FH is well documented particularly in diseases associated with excess inflammation, such as systemic lupus erythematosus and renal cell carcinomas (11-13), less is known about the importance of *S. aureus* FumC in pneumonia and how it may contribute to the inflammatory response that determines host pathology induced by *S. aureus* infection.

We hypothesize that the regulation of fumarate is an important component of *S. aureus* metabolic adaptation to the infected airway. In this report, we illustrate the unexpectedly central role of FumC and fumarate metabolism in orchestrating numerous metabolic processes in *S. aureus* relevant to its role as a pulmonary pathogen. Our findings raise the possibility of selectively targeting staphylococcal FumC to prevent chronic airway infection.

## Results

### Conservation of *fumC* across a global collection of *S. aureus* isolates

We postulated that the importance of *fumC* in *S. aureus* adaptation to the lung is reflected in its conservation across clinical isolates. Specifically, we anticipate fewer non-synonymous substitutions and protein-truncating mutations in *fumC* compared to other genes. We analyzed a global dataset comprising 820 serially collected isolates from 181 patients with cystic fibrosis (CF) and 2,139 serially collected nasal colonization isolates (570 colonization episodes) as a comparator (**Fig 1A, Tables S1-S2**). Using an established statistical genomics model of within-host evolution (11, 12), we observed a relative mutation rate, λ, less than 1 in *fumC* from CF isolates, contrasting with nasal colonizers (**Fig 1B, Tables S3-S4**). This indicates fewer mutations in *fumC* from pulmonary isolates than the mean number of mutations across the genome. Similarly, *mqo*, encoding the malate-quinone oxidoreductase enzyme involved in downstream malate metabolism to oxaloacetate (13), displayed high conservation, without any mutations in pulmonary isolates (**Fig 1B-C**). Conversely, *lqo*, which encodes a lactate-quinone oxidoreductase for lactate utilization (14), exhibited a substantially higher relative mutation rate (λ = 5.49, p = 1×10^−5^, **Table S4**).

**Figure 1.**
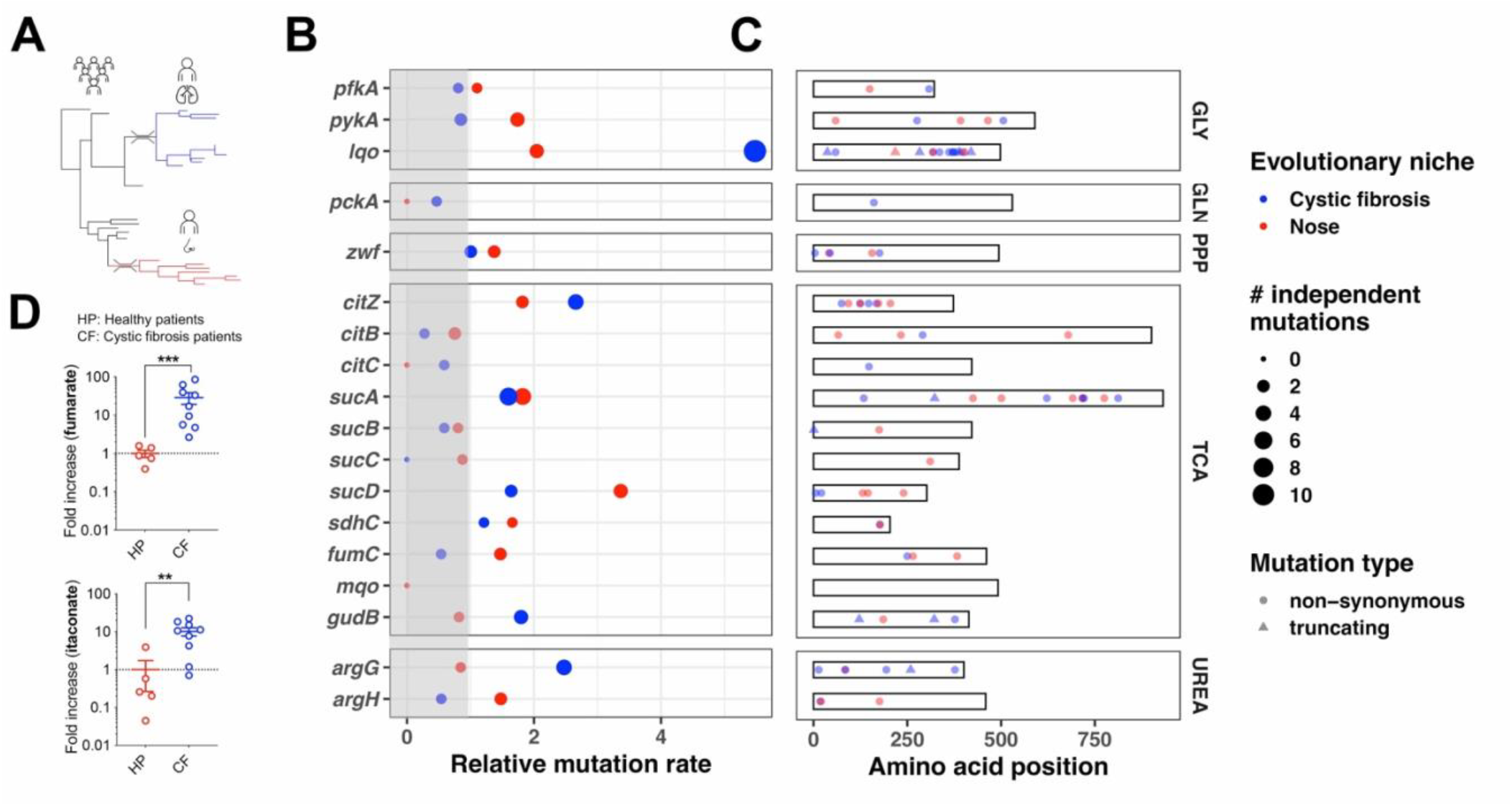
*S. aureus* adapts to the metabolic airway milieu. **(A)** Phylogenetic tree illustrating within-host evolution of *S. aureus* during persistent pulmonary infection (cystic fibrosis, blue clade) and nasal carriage (red clade). This model assumes genetic bottlenecks upon transmission and expansion of a single lineage during infection or colonization. Only mutations acquired within the host (i.e. on blue or red branches) are considered in the analysis. (**B**) Output of the convergence analysis: the size of the dots is proportional to the number of independent (i.e. acquired *de novo* within the host) protein-altering mutations in genes of the TCA cycle and surrounding pathways. The relative mutation rate is equivalent to a rate ratio in Poisson models and calculated as: (mutations in gene x/length of gene x)/(mutations in all genes/length of all genes). A rate < 1 (grey shading) indicates less mutations than the mean across the genome, suggesting that the gene is conserved during infection or colonization. GLY: glycolysis; GLN: gluconeogenesis; PPP: pentose phosphate pathway; TCA: tricarboxylic acid/Krebs cycle; UREA: urea cycle. (**C**) Gene maps with position, type, and evolutionary niche of the *de novo* mutations. (**D**) Relative levels of fumarate and itaconate in the sputum of healthy subjects (HS) and patients with cystic fibrosis (CF).

We next evaluated how the only acquired single nucleotide polymorphism (SNP), E250G, identified in *fumC* from pulmonary isolates (**Extended Data Fig 1A**) may affect the enzymatic function. Our prediction of the tetrameric structure of FumC through homology modeling and molecular dynamics (MD) simulation indicated that the glutamate residue 250 (E250) is positioned away from the active site (**Extended Data Fig 1B**), and that the SNP E250G is unlikely to inhibit its enzymatic activity. The highly negative surface charge (−78) of *S. aureus* FumC compared to the neutral human FH homologue (**Extended Data Fig 1C**) suggests differences in substrate affinity and catalytic efficiency. This observation, along with the conserved nature of *fumC* with minimal mutations in CF isolates, supports the pivotal role of FumC in chronic lung infections and its potential for selective inhibition.

### Exogenous fumarate re-directs *S. aureus* metabolic activity

We explored how high fumarate levels impact *S. aureus*. We confirmed the presence of substantially elevated levels of fumarate in the infected airways of CF patients as compared with healthy controls (**Fig 1D**). *S. aureus* preferentially utilizes glucose as a carbon source to generate energy (ATP) through glycolysis (15). In the infected lung, where the predominant recruited neutrophils compete with the bacteria for the already limited airway glucose (16), we postulate that *S. aureus* adapts to fumarate using FumC. We assessed the impact of exogenous fumarate on the transcriptional response of *S. aureu*s USA300 LAC by bulk RNA-seq analysis and observed a major impact on metabolic genes (**Fig 2A-B**). The *lacA/B/C/D/E/F* genes were significantly downregulated by fumarate (**Fig 2A-B**), indicating decreased lactose catabolism to galactose and glucose, two main carbon sources utilized by *S. aureus* for glycolysis. This was consistent with decreased glycolytic rates, which we confirmed by extracellular acidification rate (ECAR), in WT LAC and especially in the Δf*umC* mutant that cannot catabolize fumarate (**Fig 2C**). We also noted that exogenous fumarate inhibited bacterial oxidative phosphorylation (OXPHOS) represented by the oxygen consumption rate (OCR) (**Fig 2D**). The *purl/M/N/H* genes were downregulated (**Fig 2A**), suggesting decreased purine synthesis. In contrast, we observed increased expression of the *argG/H* genes, which are involved in the urea cycle and promote biofilm formation (17).

**Figure 2.**
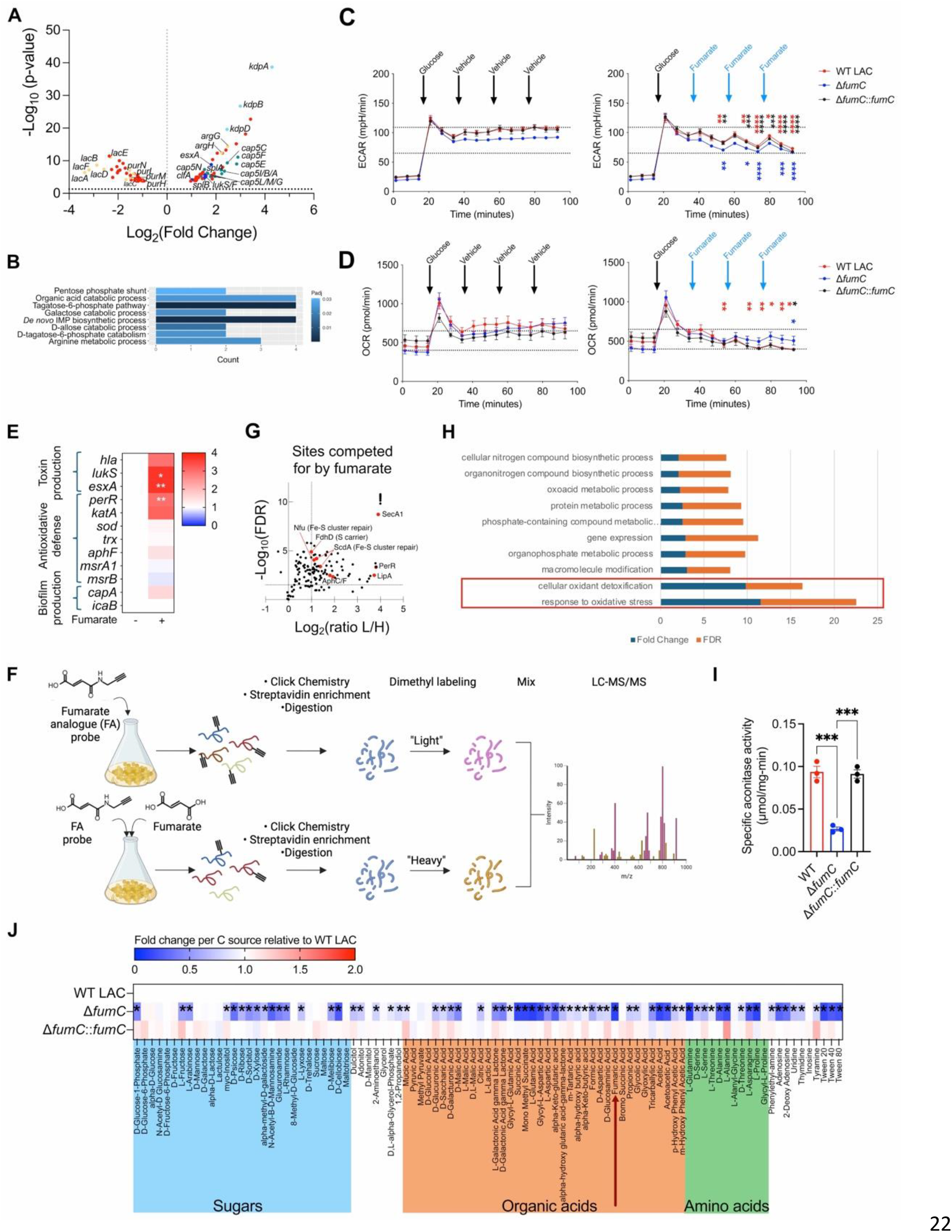
Fumarate imposes metabolic stress on *S. aureus*. **(A)** Volcano plot showing significantly differentially expressed genes in WT LAC in the presence (100 mM) and absence of fumarate above the dotted line; n = 3 biological samples (1 experiment). **(B)** Pathway enrichment analysis of genes from (A) by Gene Ontology depicting significant differences in metabolic pathways in the presence and absence of fumarate. **(C)** Glycolysis and **(D)** oxidative phosphorylation (OXPHOS) of *S. aureus*, as measured by the extracellular acidification rate (ECAR) and oxygen consumption rate (OCR) respectively, using the Seahorse extracellular flux analyzer. Glucose was injected, followed by three sequential additions of either the vehicle (left panel) or itaconate (right panel). Each data point is the mean ± SEM; n = 4 biological samples (4 independent experiments) with at least 3 technical replicates. The asterisks denote statistical differences at each time point between left and right panels by one-way ANOVA. **(E)** Heatmap illustrating variations in gene expression in WT LAC cultured with (10 mM) and without fumarate, as determined by qRT-PCR. The differences in gene expression are relative to WT LAC without fumarate. Statistical analysis was conducted using the Mann-Whitney non-parametric U test for each gene, with significance denoted as *p < 0.05, **p < 0.01, ***p < 0.001 and ****p < 0.0001. **(F)** Schematic diagram outlining the workflow for the chemoproteomic profiling of succinated *S. aureus* proteins by the fumarate analogue (FA) probe. **(G)** Volcano plot of the quantified FA-captured sites that are competed for by fumarate in live *S. aureus* FPR3757. The threshold for the -log10(q value/FDR) is set at 1.3; n = 3 biological samples (3 independent experiments) **(H)** Gene ontology analysis for the succinated sites from the *in situ* profiling. **(I)** AcnA/CitB activity in WT LAC, the Δ*fumC* mutant and complemented strain. Each data point is the mean ± SEM.; n = 3 biological samples (3 independent experiments) in triplicate. Statistical analysis was conducted by one-way ANOVA, ***p < 0.001. **(J)** Carbon source assimilation of WT LAC, the Δ*fumC* mutant and complemented strain (PM1 Biolog; n = 3 biological replicates from 3 independent experiments). The color intensity in the heatmap corresponds to the absorbance of the bacterial strains (OD_590nm_) as a readout of bacterial respiration in the presence of the indicated carbon source, normalized to the absorbance (OD_590nm_) of WT LAC in the same carbon source. Asterisks denote statistically significant differences as determined by two-way ANOVA.

The expression of genes associated with pathogenesis is often linked with bacterial metabolism (18). We observed that expression of several such genes were altered in the presence of fumarate (**Fig 2A**). The *cap5* genes associated with capsule production (19) were upregulated (**Fig 2A**). The genes *lukS* and *esxA*, encoding a PVL toxin subunit and a Type Seven Secretion System (T7SS) effector respectively, were markedly upregulated upon fumarate exposure (**Fig 2A**), as corroborated by qRT-PCR even at lower fumarate concentration (**Fig 2E**). We also observed increased expression of *perR* and *katA*, genes important in orchestrating antioxidative defenses (**Fig 2E**).

The metabolic impact of fumarate in redirecting metabolic activity can be mediated by direct or indirect mechanisms. Fumarate can post-translationally modify both bacterial and mammalian proteins, through covalent binding or succination (12, 20-22). We performed global profiling of succinated proteins in live *S. aureus* using a biorthogonal fumarate analogue (FA) probe (23) (**Fig 2F**). Among the proteins that were identified, many that were modified at specific cysteines are involved in antioxidative stress responses such as PerR (**Fig 2G-H, Table S5**). We were intrigued by targets involved in the biogenesis/repair of iron-sulfur (Fe-S) clusters, including Nfu and ScdA (**Fig 2G**). We hypothesized that fumarate impairs Fe-S cluster assembly in *S. aureus* through succination, mirroring its effect in host cells (12), and inhibits downstream Fe-dependent bacterial processes. A number of proteins require Fe-S prosthetic groups for function (24), such as the TCA cycle enzyme, aconitase (AcnA/CitB). In *S. aureus*, defects in Fe-S synthesis result in decreased TCA cycle function and reduced survival within neutrophils (25).

We tested the inhibitory effect of fumarate on *S. aureus* AcnA using the Δ*fumC* mutant as a genetic model for fumarate accumulation. We observed a significant reduction in AcnA activity in the Δ*fumC* mutant compared with the parental and complemented strains (**Fig 2I**), suggesting that fumarate accumulation may inhibit staphylococcal Fe-S assembly. Metabolite phenotype microarrays (Biolog) also highlighted the inability of the Δ*fumC* mutant to utilize numerous carbon sources, including TCA cycle intermediates and sugars (**Fig 2J**). Thus, the accumulation of excess fumarate has multiple potentially detrimental effects on the bacteria, disrupting the generation of ATP through the TCA cycle, OXPHOS, as well as glycolysis. While these studies indicate the importance of the Fe-independent enzyme FumC (26) in regulating the accumulation of fumarate, they also imply that it is involved in many bacterial biosynthetic pathways.

### Deletion of *fumC* impairs major physiological processes in *S. aureus*

We next addressed the importance of *fumC* in *S. aureus* physiology, particularly as relevant to pneumonia. We noted that under conditions that mimic the infected airway, the Δ*fumC* mutant had a significant growth defect. In artificial sputum media (ASM), which contains fumarate (**Extended Data Fig 1D**), mucin, DNA, and amino acids (27), there was a significant growth attenuation of the Δ*fumC* mutant compared to the WT and complemented strains (**Fig 3A**). This was also observed in Luria broth (LB) rich in peptides but lacking in glucose (**Fig 3B** left panel). While the WT strain and the Δ*fumC* mutant could both grow equally well in the presence of glucose in chemically defined media (CDM) (**Fig 3C** left panel), the Δ*fumC* mutant displayed significant growth impairment in the absence of glucose (**Fig 3C** right panel). This growth defect was exacerbated by the addition of fumarate and restored upon supplementation with malate in either LB, ASM or CDM (**Fig 3B** middle and right panels, **Extended Data Fig 1E**).

**Figure 3.**
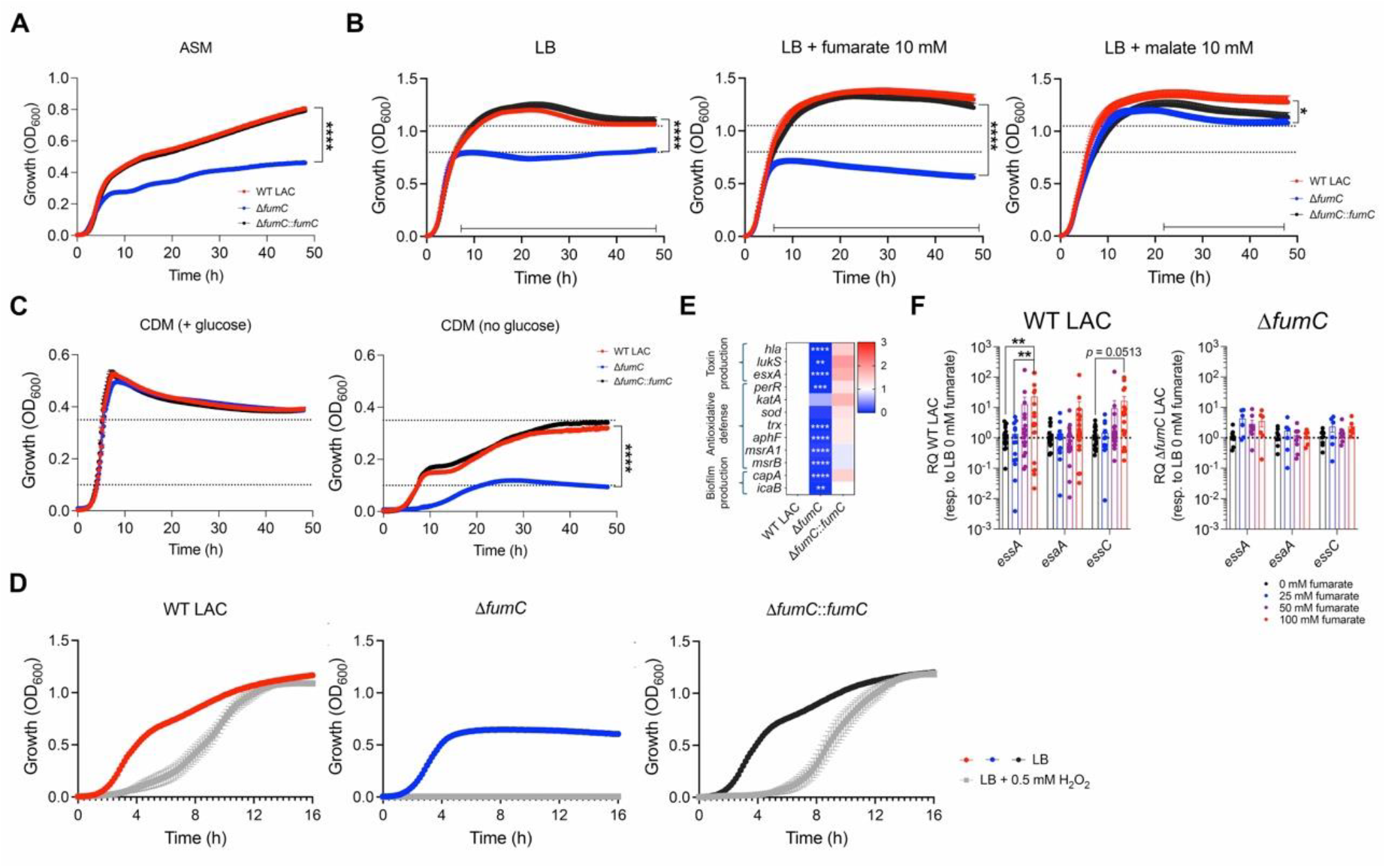
FumC promotes bacterial metabolic fitness, antioxidative defense and pathogenesis. **(A-C)** Growth curve of WT LAC, the Δ*fumC* mutant and complemented strain in **(A)** artificial sputum media (ASM), **(B)** Luria Bertani (LB) broth lacking glucose (left panel) and supplemented with fumarate (middle panel) or malate (right panel) at a final concentration of 10 mM or (**C**) chemically defined media (CDM) supplemented with (left panel) and without (middle panel) glucose. **(D)** Growth curve of WT LAC, the Δ*fumC* mutant and complemented strain in LB with and without hydrogen peroxide (H_2_O_2_) at a final concentration of 0.5 mM. Data are shown as mean ± SEM from n ≥ 3 biological replicates (≥ 3 independent experiments) at least in triplicate. Statistical significance is determined by two-tailed t-student test with FDR correction. **(E)** Heatmap showing variations in gene expression in WT LAC, the Δ*fumC* mutant and complemented strain, as determined by qRT-PCR. The differences in gene expression are relative to WT LAC. **(F)** Expression of genes encoding the T7SS machinery by WT LAC (left panel) and the Δ*fumC* mutant (right panel) in the presence of increasing fumarate concentrations. The differences in gene expression are relative to WT LAC or the Δ*fumC* mutant grown in the absence of fumarate. For **(D-E)**, statistical analyses were conducted using the two-tailed t-student test with FDR correction for each gene, with significance denoted as *p < 0.05, **p < 0.01, ***p < 0.001 and ****p < 0.0001.

In addition to effects on growth, the absence of *fumC* rendered the bacteria more susceptible to oxidative stress (**Fig 3D**), consistent with the downregulation of the antioxidative genes *perR, trx, aphF*, and *msrA1/B*, but not *katA* and *sod* that encode catalase and superoxide dismutase respectively (**Fig 3E**). Similarly, the expression of biofilm-associated genes and those encoding α-hemolysin (Hla), LukS (PVL), and the T7SS EsxA effector were significantly downregulated in the Δ*fumC* mutant (**Fig 3E**). As expected, the phenotypes associated with the Δ*fumC* mutant contrasted with those observed when *S. aureus* can degrade excess fumarate (**Fig 2E**). Of note, the expression of genes encoding the T7SS machinery itself (*essA, esaA*, and *essC)* was induced by exogenous fumarate in a *fumC*-dependent manner (**Fig 3F**). Our findings indicate that *S. aureus*, when exposed to fumarate, depends on *fumC* to express genes likely important in pathogenesis and in maintaining biosynthetic pathways crucial to preserve its physiology.

### FumC orchestrates carbon flux to diverse pathways

We elucidated the precise role of *S. aureus* FumC in the biosynthetic pathways critical for bacterial growth and fitness (**Fig 4A**), properties essential for pathogens. We employed ^13^C_4_-labeled fumarate and stable isotope tracing in WT LAC and the Δ*fumC* mutant to track the metabolic fate of fumarate. Of the total fumarate, 49.5% and 59.4% were ^13^C-labeled in WT LAC and the Δ*fumC* mutant respectively, particularly as the carbon 4 (C_4_) isotopologue (**Fig 4B**). This translated into the incorporation of ^13^C into malate as C_1_-C_4_ isotopologues in WT LAC but not in the Δ*fumC* mutant (**Fig 4B**). The accumulated ^13^C was instead incorporated back into succinate in the Δ*fumC* strain as C_3_-C_4_ isotopologues (**Fig 4B**). We observed the assimilation of ^13^C derived from labeled fumarate into the TCA cycle metabolites citrate, cis-aconitate and α-ketoglutarate (α-KG) in WT LAC but not the Δ*fumC* mutant, as predicted (**Fig 4B**).

**Figure 4.**
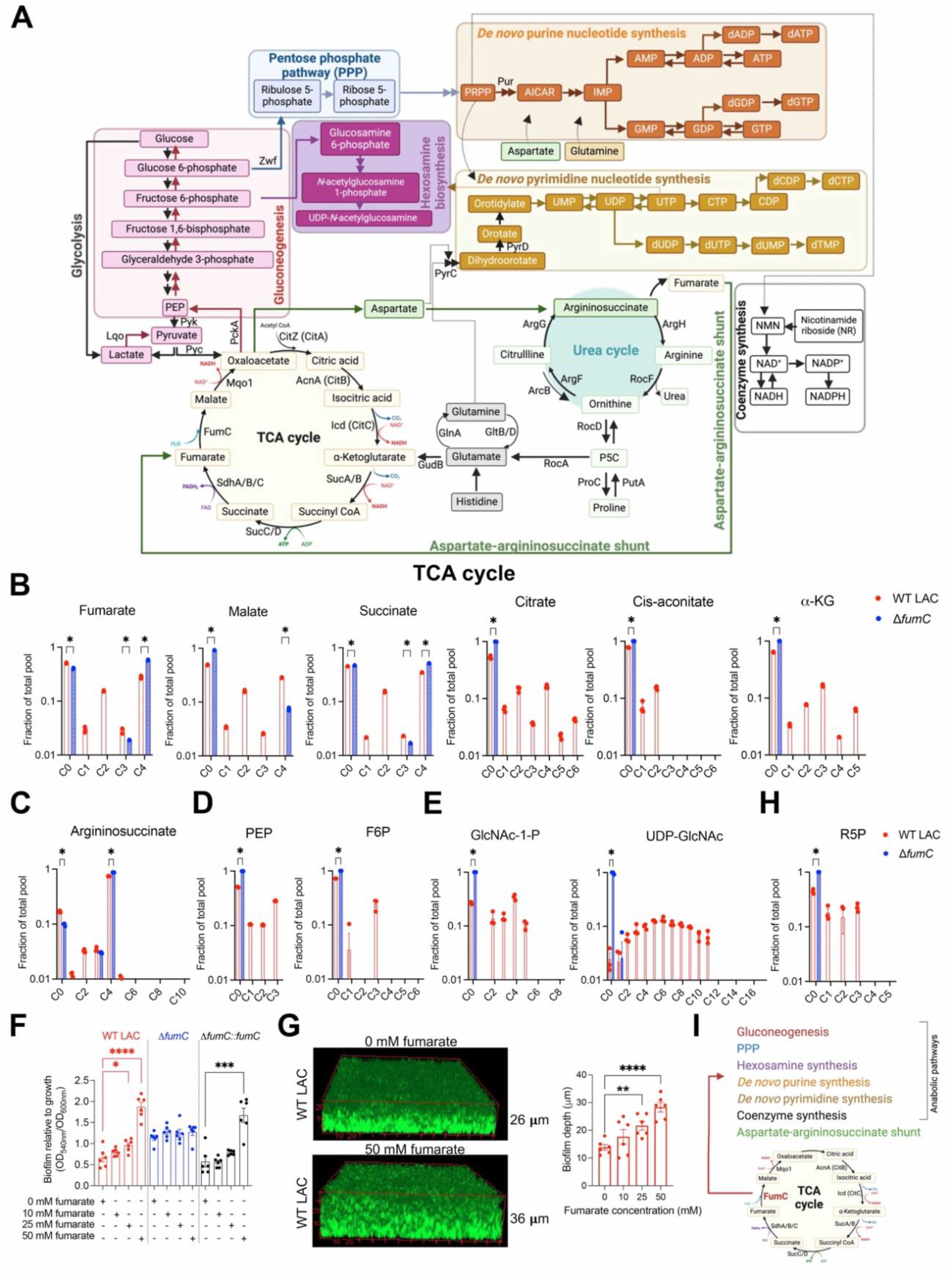
FumC serves as a metabolic hub and directs fumarate to major metabolic pathways. **(A)** Schematic diagram depicting key metabolic pathways in *S. aureus*. **(B-E, H)** ^13^C_4_-fumarate labeling of metabolites involved in the **(B)** TCA cycle or **(C)** aspartate-argininosccuinate shunt, **(D)** gluconeogenesis, **(E)** hexosamine synthesis and **(H)** PPP in WT LAC and the Δ*fumC* mutant. **(F)** Biofilm formation by WT LAC, the Δ*fumC* mutant, and the complemented strain in LB broth (without glucose) with increasing concentrations of fumarate, as evaluated by crystal violet staining. **(G)** Biofilm depth of WT LAC in the presence of increasing fumarate concentrations, measured using wheat germ agglutinin-Alexa Fluor 555 staining and confocal microscopy. For **(B-H)**, data are shown as mean ± SEM from n = 3 biological replicates (1 independent experiment), or n = 2 biological replicates for **(G)**. Statistical significance is determined by two-tailed t-student test with FDR correction. **(I)** Diagram summarizing the anabolic pathways fueled by *S. aureus* FumC.

We detected the substantial integration of labeled carbon from fumarate into argininosuccinate, particularly as the C_4_ isotopologue (**Fig 4C**). This may serve to rapidly replenish malate and oxaloacetate for gluconeogenesis, hexosamine synthesis and the pentose phosphate pathway (PPP) in the WT strain (**Fig 4A**). Accordingly, there was increased integration of ^13^C into the gluconeogenic intermediates, phosphoenol pyruvate (PEP) and fructose 6-phosphate (F6P), in WT LAC but not in the Δ*fumC* mutant (**Fig 4D**). We also detected significant incorporation of ^13^C into *N*-acetyl-glucosamine-1-phosphate (GlcNAc-1-P) and UDP-*N*-acetyl-glucosamine (UDP-GlcNAc) (**Fig 4E**), which contribute to biofilm formation (9, 28) in the WT strain. We confirmed that fumarate promoted biofilm formation in a dose-dependent manner in WT LAC (**Fig 4F**), resulting in increased biofilm depth (**Fig 4G**).

In addition, we noted the incorporation of labeled carbon into the PPP intermediate, ribose 5-phosphate (R5P) in WT LAC (**Fig 4H**). This was not observed in the Δ*fumC* mutant. Assimilation of ^13^C into R5P correlated with the increased labeling of intermediates in the *de novo* purine nucleotide metabolism, such as AICAR, AMP, ADP, ATP, GMP and GDP (**Extended Data Fig 1F**). In parallel, there was increased labeling of dihydroorate, orotate and intermediates of the *de novo* pyrimidine nucleotide synthesis, including UMP, UDP, UTP and CTP, in WT LAC only (**Extended Data Fig 1G**). Consistently, the incorporation of ^13^C into NAD and the reduced form NADH, which is crucial for energy production, occurs in the WT strain only (**Extended Data Fig 1H**). Overall, the stable isotope tracing experiment emphasized an unexpectedly central role of FumC as a metabolic hub, directing fumarate utilization into multiple biosynthetic pathways critical for *S. aureus* survival and proliferation (**Fig 4I**).

### FumC is critical in pneumonia

We expected *S. aureus* FumC to be crucial in chronic lung infection. We used a mouse pneumonia model with elevated fumarate levels to more adequately mimic the airway milieu in persistent infections (**Fig 1D**). *Ptenl*^*-/-*^ mice, which were previously used to model the airway properties in CF (29-31), exhibited a baseline accumulation of the pro-inflammatory metabolite fumarate in the lung (approximately 5 times higher than in BL/6 mice) but not succinate (**Fig 5A**). As predicted, in settings with high fumarate levels (*Ptenl*^*-/-*^), we observed that the Δ*fumC* mutant was significantly attenuated compared to the WT strain (**Fig 5B**). This attenuation was more modest in the bronchoalveolar lavage fluid (BALF) and lung of BL/6 mice (3-fold and 5-fold respectively) (**Fig 5B**). Deletion of *sucD*, encoding the TCA cycle enzyme succinyl-CoA synthetase, had no effect on airway *S. aureus* load, while the Δ*sucA* mutant, lacking α-KG dehydrogenase, was attenuated in the lung but not in the BALF (**Extended Data Fig 1I**). This suggests that not all TCA cycle disruptions negatively impact staphylococcal airway survival and highlights the importance of fumarate metabolism in pulmonary pathogenesis.

**Figure 5.**
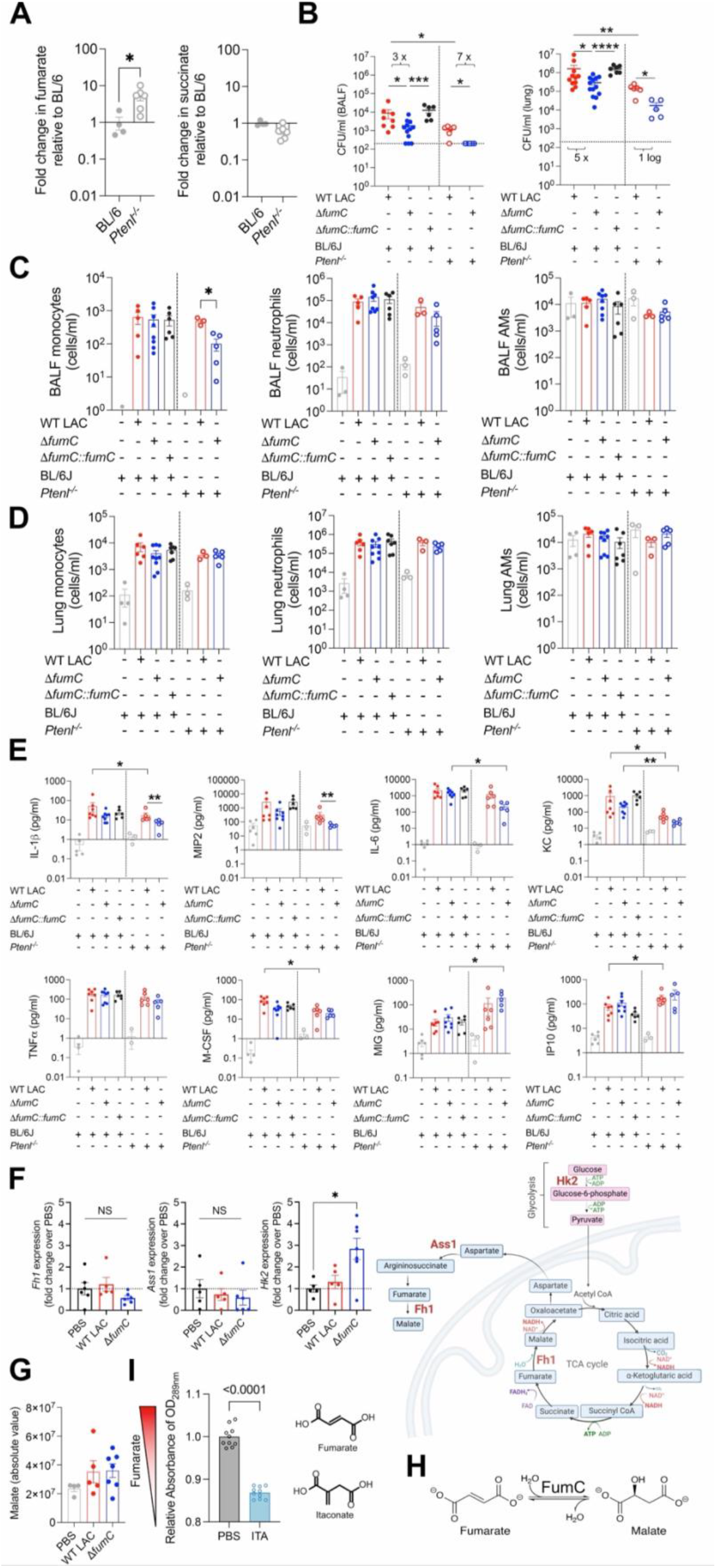
Host fumarate hydratase and *S. aureus* FumC both regulate fumarate and pathogenesis. **(A)** Relative level of fumarate (left panel) and succinate (right panel) in the BALF of uninfected BL/6 and *Ptenl*^*-/-*^ mice. **(B)** Bacterial burden from the BALF (left panel) and lung (right panel) of BL/6 and *Ptenl*^*-/-*^ mice infected with WT LAC, the Δ*fumC* mutant and complemented strain; BL/6 infected with WT LAC (*n* = 11), Δ*fumC* mutant (n = 13), and Δ*fumC*::*fumC* (n = 7), *Ptenl*^*-/-*^ infected with WT LAC (n = 6), Δ*fumC* mutant (n = 5). The above-mentioned total number of mice per group were from at least 2 independent experiments The dotted line indicates the limit of detection. **(C-D)** Innate immune cells (monocytes left panel, neutrophils middle panel and alveolar macrophages right panel) from the **(C)** BALF and **(D)** lungs of uninfected and infected mice from (A). **(E)** Cytokine measurements from the BALF of uninfected and infected mice from (A). **(F)** Expression of murine *Fh1, Ass1* and *Hk2* from uninfected and infected mouse lungs (n = 5 per condition) by qRT-PCR (left panel) and schematic diagram showing the metabolic reactions catalyzed by the encoded enzymes fumarate hydratase, argininosuccinate synthase and hexokinase (right panel). **(G)** Absolute level of malate in the BALF of uninfected (PBS) BL/6 mice and mice infected with WT LAC or the Δ*fumC* mutant. **(H)** Schematic diagram of reaction catalyzed by FumC. **(I)** FumC activity assessed by the absorbance of fumarate (left panel). Fumarate was supplied as a substrate for FumC, preincubated with and without 1 mM itaconate (chemical structure, right panel). Data represent mean values ± SEM. Statistical analysis was performed by Mann-Whitney non-parametric U test, *p < 0.05, **p < 0.01, ***p < 0.001 and ****p < 0.0001.

To determine if differences in the immune response due to the loss of *Ptenl* might account for the enhanced Δ*fumC* clearance, we compared the immune cells (neutrophils, monocytes, alveolar macrophages) and cytokines during infection of both BL/6 and *Ptenl*^*-/-*^ mice. Overall, similar numbers of immune cells were observed in the infected BL/6 and *Ptenl*^*-/-*^ airway (**Fig 5C-D**). In the *Ptenl*^*-/-*^ mice, where the difference in infection levels between WT LAC and the Δ*fumC* mutant was more pronounced (**Fig 5B**), we observed a slight reduction in monocyte numbers in the BALF but not in the lung (**Fig 5C-D**). While these mice also had lower levels of IL-1β and MIP2, there were no statistical differences in other proinflammatory cytokines such as IL6, KC and TNFα (**Fig 5E**). In the BL/6 mice, similar numbers of immune cells and cytokine levels were also detected in response to infection with either the WT or Δ*fumC* strains (**Fig 5C-E**). We confirmed that there were no differences in the uptake of WT LAC and the Δ*fumC* mutant by phagocytes (**Extended Data Fig 1J**). Thus, our results indicate that the attenuation of the Δ*fumC* mutant is not due to a lack of immunostimulation or to differences in the immune response between BL/6 and *Ptenl*^*-/-*^ backgrounds. Instead, our findings underscore the critical role of *S. aureus* fumarate metabolism during pneumonia.

### *S. aureus* exploits host metabolism

The apparent disparity observed between the profound *in vitro* effects of *fumC* on *S. aureus* metabolism and the modest attenuation of the Δ*fumC* mutant in BL/6 mice led us to speculate that the bacteria may exploit the host FH to mitigate the absence of *fumC in vivo*. We observed comparable *Fh1* transcript levels in both uninfected and infected murine lungs (**Fig 5F**). This indicates that *S. aureus* sustains host *Fh1* expression, in contrast to Gram-negative pathogens and LPS, which typically downregulate it (11, 32). The expression of *Ass1*, which encodes argininosuccinate synthase 1, was also unchanged, unlike *Hk2* that was significantly upregulated upon infection with the Δ*fumC* mutant (**Fig 5F**). We next measured the level of malate, the product of either *S. aureus* FumC or host FH, in the BALF harvested from BL/6 mice infected with WT LAC or the Δ*fumC* mutant. Similar levels of airway malate were detected during infection with either strain (**Fig 5G-H**) despite the inability of the Δ*fumC* mutant to catabolize fumarate to malate. This suggests that host FH activity may protect S. *aureus* from the detrimental effects of fumarate *in vivo*, thereby limiting the impact of *fumC* deletion through the generation of malate. Our results emphasize that while *S. aureus* can exploit the host FH to degrade fumarate, it depends significantly on its own FumC enzyme in environments with accumulated fumarate levels typical of chronic lung infections.

Itaconate, another dicarboxylate, accumulates in the chronically infected airway, along with fumarate (**Fig 1D**). These two metabolites share structural similarities (**Fig 5I**) and could potentially work in a concerted fashion. We wondered if the presence of itaconate might further enhance FumC activity. We monitored the conversion of fumarate to malate by purified FumC (33) in the presence or absence of itaconate (**Fig 5H-I**). The significant depletion of added fumarate in the presence of itaconate strongly suggests that itaconate potentiates FumC activity. Our data demonstrate that the metabolism of airway fumarate by S. *aureus* promotes pneumonia and that this effect is even more biologically significant in the context of other accumulated metabolites such as itaconate.

## Discussion

*S. aureus* is a well-studied pathogen which nonetheless remains among the top three causes of healthcare-associated infection, especially pneumonia (1). Its extensive metabolic flexibility has long been recognized but not effectively targeted in either antibiotic or vaccine development. We uncovered a previously unappreciated role for fumarate metabolism in the overall physiology and pathogenesis of *S. aureus*, especially in microenvironments with limited glucose such as the airway. Our focus on FumC was initiated by studying clinical isolates that had successfully undergone long term adaptation to the lung. The increased expression (10) and low mutation rate of *fumC* in *S. aureus* pneumonia isolates indicate its global epidemiological significance for pulmonary adaptation. This is particularly relevant in the chronically infected airway replete with the host immunometabolites, fumarate and itaconate.

We observed a fascinating dynamic regulation of airway fumarate, an important pro-inflammatory mediator (11), by both host and *S. aureus* fumarate hydratase. For the host, the production of the dicarboxylates fumarate and itaconate offers homeostatic protection against the detrimental effects of excessive inflammation. Fumarate, a product of activated macrophages, induces type I interferon and TNFα production in response to mitochondrial damage (11). In contrast, itaconate exhibits immunomodulatory properties; it stimulates major antioxidative mediators such as Nrf2 and ATF3 (34) and inhibits NLRP3 inflammasome activation (35). Just as excess fumarate is detrimental to the host, we show that it is toxic to the bacteria, inhibiting primary metabolic pathways such as glycolysis, the TCA cycle and OXPHOS. The presence of *fumC* in *S. aureus* isolates not only enables fumarate degradation, circumventing its negative effects, but also promotes its utilization in anabolic pathways such as gluconeogenesis, and hexosamine production, which are essential for biofilm formation and defense against oxidative stress. We also found that several genes expected to function more directly in pathogenesis were upregulated by fumarate such as those involved in toxin and capsule production and the T7SS.

Having documented the pervasive effects of fumarate on *S. aureus* biology and the critical role of FumC in numerous metabolic pathways, we also demonstrate its role *in vivo*, particularly in environments abundant in fumarate, such as the *Ptenl*^*-/-*^ airway. Consistently, we show that the Δ*fumC* mutant had a more modest limitation during acute pulmonary infection in BL/6 mice. No substantive differences in the immune response to the WT or Δ*fumC* strains were identified. These observations can be explained by compensatory host FH activity, as confirmed by quantifying equivalent levels of malate in the airway of BL/6 mice. Thus, the host can regulate local fumarate levels to a certain extent, even in the absence of *S. aureus* FumC, enabling the microorganisms to proliferate. However, in the *Ptenl*^*-/-*^ background with excessive airway fumarate, the lack of *fumC* resulted in an appreciable loss of bacterial fitness *in vivo*. This further supports the critical role of FumC in chronic pulmonary infections.

In addition, we observed that the host metabolite itaconate enhances the activity of *S. aureus* FumC and further promotes the degradation of fumarate and its utilization as a carbon source. Thus, *in vivo*, ambient fumarate amounts are regulated by both the host FH and *S. aureus* FumC whose activity seems likely to be boosted by itaconate. It remains to be determined exactly how itaconate accelerates FumC and whether this is unique to *S. aureus*. Given the overall structural similarities between the host and *S. aureus* FH, FumC may initially appear to be an unsuitable target for anti-staphylococcal therapy. However, some of the peculiarities of *S. aureus* FumC structure and charge characteristics may be sufficiently distinct to generate selective bacterial toxicity. These distinctive features could be harnessed for the development of adjuvant therapy to prevent chronic pneumonia.

## Methods

### *In silico fumC* conservation

We assembled a collection of serially sequenced *S. aureus* strains from seven published within-host evolution studies of *S. aureus* during cystic fibrosis (CF) (36-42). The studies were added if: they included at least two *S. aureus* strains per patient; sequencing reads were publicly available; minimal metadata were reported: patient identifier, specimen source, date of collection. We supplemented the published studies with unpublished data from a CF study at the Children’s Hospital of Philadelphia and with seven published studies of *S. aureus* nasal colonization (43-49). Raw reads and metadata were downloaded from NCBI using the Bioproject identifier provided in the publications (**Table S2**). Metadata were additionally retrieved from the publications. Quality control for reads data was performed as described in (50). Reads were excluded if the coverage was below 20, reads classification using Kraken2, v2.1.1 (51) did not yield *S. aureus* or <50% reads were classified as *S. aureus*, or if the assembly size exceeded 3.5 million bp. Reads were assembled using Shovill, v.1.1.0, which implements the Spades assembler (52). Multi-locus sequence type and resistance genes were inferred from the assemblies using Mlst, v2.19.0 and Abricate, v1.0.1, both available at https://github.com/tseemann.

Within-host evolution analysis was carried out as described in (53): for each patient or nasal carrier we used Snippy, v4.6.0 (https://github.com/tseemann/snippy) to map isolate reads to the draft assembly of the earliest strain (if multiple strains were available the reference was selected randomly). The output was filtered by removing variants from reference reads and variants at positions for which the reads coverage of the reference was below 10. The amino-acid sequences of the mutated genes across all episodes were clustered using CD-HIT, v4.8.1 (54) and for each cluster representative we searched for a *S. aureus* FPR3757 (55) homolog (NCBI accession CP000255) using Blastp, v2.10.1. This generated a list of FPR3757 genes with counts of independent episodes with at least one mutation arising within the host. We then excluded synonymous substitutions and limited the list to genes involved in the tricarboxylic acid cycle (TCA) and surrounding pathways (glycolysis, gluconeogenesis, urea cycle) and calculated the relative mutation rate λ for each gene i as (mutations in gene i/length of gene i)/(mutations in all genes/length of all genes). We inferred the significance of the mutations enrichment by fitting two Poisson models: model 0 as the null hypothesis (neutral evolution), where the number of variants per gene was the product of the mean mutation rate across the genome and the gene length; model 1 estimated the number of variants per gene base on a gene-specific mutation rate while variants in the remaining genes are the product of a mean mutation rate. Model 0 and model 1 were compared using a likelihood ratio test. Scripts for the within-host evolution analysis are available at https://github.com/stefanogg.

### Bacterial strains

The bacterial strains used in this study are shown in Table S6. WT LAC and its derivative strains, Δ*fumC* and Δ*fumC::fumC* were constructed as previously described (56). The transposon insertions *sucA::Tn*(*ermB*) and *sucD::Tn*(*ermB*) were sourced from the NARSA transposon mutant library (57). The alleles were transduced into USA300_LAC using bacteriophage 80α (58). The mutant strains were confirmed via PCR using sucAB and sucCD primers. All strains were grown at 37°C on plates of Luria-Bertani lacking glucose (LB, Becton Dickinson (BD), 244610) supplemented with 1% agar (w/v, Acros Organics, 400400010). Overnight cultures and subcultures (1/100) were grown in LB broth at 37°C with shaking. Bacterial inocula were estimated based on the optical density at 600nm, OD_600nm_, and verified by retrospective plating on LB agar plates to determine colony forming units (CFUs). *S. aureus* recovered from the mouse airways were selected on LB agar plates and BBL CHROMagar Staph aureus plates (BD).

### Primers

The primers used in this study are listed in Table S7. Full-length *fumC* was amplified from the genome of WT USA300 FPR3757 using KOD One PCR Master Mix -Blue- (TOYOBO, KMM-201) and cloned into the pET-21b(+) vector by Gibson assembly.

### Growth curves

A U-bottomed, clear 96-well plate (Greiner Bio-One #650161) was prepared with LB, chemically defined media (CDM) (59) or artificial sputum media (27). For the experiments that used media supplemented with itaconic acid (Sigma, I29204), L-malic acid (Sigma, M1000), sodium fumarate (Sigma, F1506), or hydrogen peroxide, H_2_O_2_, (Sigma, H1009), the pH of the media was corrected to 7.0 prior to filter sterilization using 0.20 °m filters (Thermo Fisher Scientific,725-2520). Each well was inoculated with 1.5 × 10^5^ bacteria. Absorbance at 600 nm was read every 10 minutes for 36 h on the Varioskan™ LUX multimode microplate (Thermo Fisher Scientific, VL0000D0), as the plate was maintained at 37°C with shaking.

### Measurement of biofilms

Log phase cultures were diluted to OD_600nm_ of 0.1 in LB broth supplemented with or without fumarate (Sigma, F1506) prior to incubation in either 96-well plates (Greiner Bio-One, 650161, 100 μl/well) or 4-well chamber slides (Sarstedt, D-51588, 750 μl/well) for 24 h and 12 h respectively. The plates were processed as follows:

1. Crystal violet staining in 96 well-plates: The OD_600nm_ was measured prior to crystal violet staining. The plates were fixed with 100% methanol, stained with 1% crystal violet (w/v, Sigma, C6158), washed and dried. Subsequently, 33% acetic acid (v/v, Acros Organics, 222140010) was added to solubilize the dye. The OD_540nm_ was measured and normalized to the bacterial growth at OD_600nm_.
2. Wheat germ agglutinin-Alexa Fluor 555 staining and confocal microscopy: Wheat germ agglutinin (WGA) coupled to Alexa Fluor 555 (5 μg/ml, Thermofisher, W32464) was added into each well and the chamber slides were covered with foil. Each chamber slide was placed on a shaker (horizontal orbit, 60 rpm) in a 37°C room for 5 minutes, before being kept stationary in a 37°C room for 12 hours. Confocal imaging of viable biofilms was performed using a Zeiss LSM 980 Airyscan with a 37°C cage incubator. 16-bit images of WGA-positive cells (1.5x crop area, 1024×1024 px frame size) were captured using a 40X oil-immersion objective with the 488 nm laser (670 V gain) set at 1% intensity. Z-stack slices were collected at 1 μm intervals. The z-depth of high-density biofilm layers were calculated as the number of slices (1 μm apart) where extracellular spaces occupied no more than 20% of the image. ImageJ 1.54f. (National Institutes of Health, Bethesda, MD, USA) was used to calculate the % of extracellular space in an image using thresholding.

### Isolation of RNA from bacterial cultures

Bacterial cultures grown in LB with and without 10 mM fumarate were standardized to an OD_600nm_ of 1.0 prior to centrifugation. Bacterial pellets were incubated in lysis buffer (50 μM Tris-EDTA pH 7.5, 8 U ml^−1^ mutanolysin (Sigma-Aldrich), 0.018 mg ml^−1^ lysostaphin (Sigma-Aldrich), 0.05 mg ml^−1^ lysozyme (Sigma-Aldrich)) at 37 °C for 45 min and TRK lysis buffer (Omega Bio-tek) was added. After 10 min at room temperature, 70% ethanol was added, and samples were transferred to E.Z.N.A. RNA isolation columns (Omega Bio-tek). RNA was isolated following the manufacturer’s instructions and treated with DNase using the DNA-free DNA removal kit (Invitrogen).

### Complementary DNA synthesis and qRT-PCR

Multiscribe reverse transcriptase (Applied Biosystems) was used to generate cDNA for qRT–PCRs with Power SYBR Green PCR Mastermix (Applied Biosystems). qRT-PCR was performed using primers for *hla, lukS, esxA, per, katA, sod, trx, aphF, msrA1, msrB, capA, icaB and 16S* (Table S7) and a StepOne Plus thermal cycler (Applied Biosystems). The data were analyzed using the ΔΔC_T_ method.

### RNA-sequencing

WT LAC was grown in LB with or without 100 mM fumarate to late exponential phase. Bacterial pellets were incubated in a cell wall lysis mixture (described above) at 37°C for 45 min. TRK lysis buffer (Omega Bio-tek, R6834-02) and 70% ethanol were added, and samples were transferred to E.Z.N.A. RNA isolation columns (Omega Bio-tek, R6834-02). RNA was isolated following the manufacturer’s instructions and treated with DNase using the DNAfree DNA removal kit (Fisher Scientific, AM1906). The RNA was precipitated with 0.1 volume 3 M sodium acetate (Thermo Fisher, S209) and 3 volumes of 100% ethanol, recovered by centrifugation, and washed with ice cold 70% ethanol. A ribosomal RNA-depleted cDNA library was prepared according to the manufacturer’s instructions using the Universal Prokaryotic RNA-Seq Prokaryotic AnyDeplete kit (NuGEN, 0363-32) and sequenced with Illumina HiSeq. Raw base calls were converted to fatsq files using Bcl2fastqs. Filtered reads were aligned to the LAC_FPR3757 reference genome using STAR-Aligner v2.7.3a. The mapped reads were annotated for read groups and marked for duplicates using the Picard tools v2.22.3. The raw counts were quantified using Subreads:FeatureCounts v1.6.3 and processed for differential gene expression using DEseq2 in R v3.5.3.

### *In silico* protein structure simulation and charge prediction

The predicted structure of *S. aureus* FumC was constructed using the homology modeling tool from the Schrödinger software package (Schrödinger Release 2022-4: Prime; Schrödinger, Inc: New York, NY, 2022) (60). We used the human fumarate hydratase (PDB: 5UPP) with 57% identity as template. The structure of *E. coli* FumC, which shares a higher identity (60%), could have been utilized, but would have necessitated the *de novo* construction of longer loops; one loop consisting of 3 amino acids and another loop consisting of 8 amino acids instead of two loops of 3 amino acids and one loop of 1 amino acid. The obtained structure was replicated into four copies, which were superimposed on the four chains of the template tetramer structure and merged. Protonation states of the predicted tetramer were assigned using PROPKA and the structure subjected to a constrained minimization using the OPLS4 force field (Schrödinger Release 2022-4: Protein Preparation Wizard, Schrödinger, LLC, New York, NY, 2022) (61).

### Synthesis of FA-alkyne probe

Monomethyl fumarate (1 g, 1 eq, 7.69 mmol) was dissolved in acetonitrile and cooled to 0°C. EDCl·HCl (1.6 g, 1.1 eq, 8.46 mmol), HOBt (1.1 g, 1.1 eq, 8.46 mmol) and Et_3_N (1.39 mL, 1.3 eq, 2.29 mmol) were added and then stirred for 40 min at 0°C. Subsequently, propargylamine (0.6 g, 1.3 eq, 10 mmol) was added. The solution was allowed to warm up to room temperature and stirred overnight. The solvent was removed *in vacuo*, the residue was redissolved in ethylacetate and washed with 1 M HCl, saturated sodium bicarbonate solution and brine. The organic layer was dried over anhydrous sodium sulfate, the resulting crude was dried, and the solvent was removed *in vacuo*. The resulting crude was purified by flash column chromatography (SiO_2_, DCM:MeOH=40:1) to afford the pure compound **1** (820.8 mg, 63.9%) as white solid (**Extended Data Figure 2A**). Lithium hydroxide (330.0 mg, 7.86 mmol) was added to a stirring solution of compound **1** (820.8 mg, 4.91 mmol) in a 1:1 mixture of water and tetrahydrofuran (**Extended Data Figure 2B**). The reaction mixture was stirred at room temperature and monitored by TLC. After 1 h, the reaction mixture was acidified with 1M HCl and the solvent was removed *in vacuo*. The crude product was redissolved in ethylacetate and filtered. The solution was evaporated *in vacuo* and the crude product was purified by flash column chromatography (SiO_2_, DCM:MeOH=20:1 to 10:1) to obtain the pure FAyne (FA-alkyne, 622.1 mg, 82.8%) as white solid. ^1^H NMR (400 MHz, DMSO-*d*_*6*_) δ 8.96 (t, J = 5.5 Hz, 1H), 6.94 (d, J = 15.5 Hz, 1H), 6.58 (d, J = 15.5 Hz, 1H), 4.01 (dd, J = 5.5, 2.6 Hz, 2H), 3.21 (t, J = 2.5 Hz, 1H) (**Extended Data Figure 2C**). ^13^C NMR (101 MHz, DMSO-*d*_*6*_) δ 166.28, 163.35, 135.86, 131.37, 81.77, 72.78, 28.64 (**Extended Data Figure 2D**).

### Competitive ABPP workflow for *S*-succinated cysteines in *S. aureus*

*S. aureus* USA300 FPR3757 was cultured overnight in tryptic soy broth (TSB), washed with PBS and concentrated to OD_600nm_ of 40. For the competition group, 800 μl of bacterial suspension was incubated with 80 μl of fumarate (100 mM stock was dissolved in NaOH and pH was adjusted to neutral) on the ThermoMixer (950 rpm, 30 min, 37°C). For the control group, fumarate was replaced with the same volume of PBS. After competition, both groups were incubated with 8 μl of 1 M FA-alkyne probe on the ThermoMixer (950 rpm, 37°C, 1 h). The bacterial suspension was then centrifuged (12,000 rpm, 5 min, 4°C), the supernatant was decanted, and the pellets were washed once with 1 ml of pre-chilled PBS. Washed pellets were resuspended with 200 μl of 0.1% (v/v) Triton X-100 (Amresco, 0694-1L) in PBS (hereafter, 0.1% PBST). Bacterial suspension was incubated on the ThermoMixer (1,200 rpm, 15 min, 37 °C) after the addition of 3 μl of lysostaphin (5 U/μl, Coolaber, CL6941-1mg). Then, the lysate was supplied with 6 μl of 10% SDS (Thermo Fisher Scientific, AM9822) and sonicated (35% intensity). The lysate was centrifuged (20,000 g, 10 min, RT) to remove the debris and the supernatant was transferred into a new 1.5 ml tube (30 μl of supernatant was also aliquoted for the gel-based ABPP assay). Click reaction was carried out with 106 μl of Click reagent mix (60 μl of 50 mM TBTA ligand, 20 μl of 50 mM CuSO_4_, 20 μl of freshly prepared 50 mM TCEP and 6 μl of 20 mM acid-cleavable azide-biotin tag (Confluore, BBBD-19-25mg) and incubated on the ThermoMixer (1,200 rpm, 1 h, 29°C). The samples were then subjected to streptavidin enrichment and dual enzyme digestion (trypsin and Glu-C) steps before mass spectrometry analysis as previously described (62).

After the dual enzyme digestion, the peptides were further labeled with a dimethylation reagent (“light” for the control group (i.e., FA probe + vehicle) and “heavy” for the fumarate competition group (i.e., FA probe + fumarate)) (63). Streptavidin beads were washed three times with MS-grade water to thoroughly remove dimethylation reagents. The labeled peptides were released from streptavidin beads by incubating with 200 μl of 2% formic acid on the ThermoMixer (1,200 rpm, 25°C, 90 min). Peptides from the control group and fumarate competition group were combined and dried in a SpeedVac (30°C). They were then analyzed by HPLC-MS/MS and subjected to quantitative proteomic analysis. A competitive light-to-heavy ratio for each target was calculated. A ratio greater than 1 (or greater than 0 after log2 transformation) suggests that the cysteine site (or protein) undergoes succination by fumarate. The corresponding competition ratios for each cysteine site (or protein) were averaged across three biological replicates, and the q-value (false discovery rate/FDR) was calculated.

### Protein purification

Plasmid pET21b-FumC-His_6_ was transformed into BL21(DE3). Protein expression was performed overnight at 16°C in the presence of 0.2 mM IPTG (VWR, 0487-100G). Bacterial pellets were resuspended in suspension buffer (50 mM Tris-HCl, pH 8.0 (Thermo Fisher Scientific, 15568-025), 150 mM NaCl) and disrupted with sonication. After the lysates were clarified via centrifugation (12,000 rpm, 30 min, 4°C), the supernatant was loaded onto 5 ml HisSep Ni-NTA 6FF column (YEASEN, 20504ES25) and washed with W1 (50 mM Tris, pH 8.0, 150 mM NaCl, 20 mM imidazole, pH 7.4) and W2 (50 mM Tris, pH 8.0, 150 mM NaCl, 40 mM imidazole, pH 7.4). Protein was eluted with elution buffer (50 mM Tris, pH 8.0, 150 mM NaCl, 300 mM imidazole, pH 7.4) and the eluate was centrifuged (4,000 g, 4 °C, swing-out) using Amicon® Ultra-15 Centrifugal Filter Unit to remove imidazole and concentrate protein. The concentrated protein was mixed with 20% glycerol and subjected to BCA assay to measure concentration (Thermo Fisher Scientific, 23225). Finally, the protein was supplied with 1 mM DTT, aliquoted, flash frozen in liquid nitrogen, and stored at −80°C.

### FumC activity

Recombinant FumC was diluted with Tris-HCl (20 mM, pH 8.0) to 1 mg/ml, and incubated with either 1 mM itaconate (pH was adjusted with NaOH to 7.4) and 1 mM DTT or PBS and 1 mM DTT as the negative control (2 h, 37°C). For the enzymatic assay, the reaction was conducted in a 1.5 ml polystyrene cuvette (BIOFIL, CUV010015). The reaction was initiated by mixing 100 μg of post-incubated FumC and the reaction mixture containing 20 mM Tris-HCl (pH 8.0) and 15 mM fumarate (Sigma-Aldrich, 47910) in a total volume of 1 ml at 25°C for 30 min. The maximal absorbance of fumarate was determined using a spectrophotometer (IMPLEN, P330) in a 1.5 ml cuvette. Absorbance was read at 289 nm. This enzymatic assay was performed on three biological replicates.

### AcnA/CitB activity

AcnA assays were conducted as previously described with slight modifications (64, 65). Strains were cultured overnight in TSB before diluting them in fresh TSB to an OD_600nm_ of 0.1. The cultures were diluted in 0.5 ml or 4 ml of TSB in 10 ml culture tubes. The cells were cultured for 8 h, before they were harvested by centrifugation, and the cell pellets were stored at −80°C. The cells were thawed anaerobically, resuspended with 200 μl of AcnA buffer (50 mM Tris, 150 mM NaCl, pH 7.4), and lysed by the addition of 4 μg lysostaphin and 8 μg DNase. The cells were incubated at 37°C until full lysis was observed (∼1 h). The cell debris was removed by centrifugation, and AcnA activity was assessed by monitoring the conversion of isocitrate to cis-aconitate spectrophotometrically.

### Metabolite phenotype microarray (Biolog)

For Carbon Source Phenotype Microarray™ (Biolog) assays, a stock solution of 2 × 10^7^ bacteria/ml was prepared in 1X IF-Oa buffer (Biolog, 72268) supplemented with 1X Redox Dye Mix A (Biolog, 74221). 100 *μ*l of this stock solution (delivering 2 × 10^6^ bacteria) was added to each well of a PM1 Microplate™ (Biolog, 12111) and the plate was incubated at 37°C overnight with shaking. Absorbance was read at 590 nm on the SpectraMax M2 plate reader (Molecular Devices).

### Bacterial extracellular flux analyses

The XFe24 sensor cartridge (Agilent #102340-100) was calibrated as per the manufacturer’s instructions overnight at 37°C without CO_2_. 500 μl of XF base medium (Agilent #102353-100) was added to each well of a Seahorse XF24 well plate (Agilent #102340-100) and inoculated with 3 × 10^7^ bacteria for a 3 h incubation at 37°C. The oxygen consumption rate (OCR, indicative of oxidative phosphorylation) and extracellular acidification rate (ECAR, indicative of glycolysis) were again measured on a Seahorse XFe24 Analyzer (Agilent Technologies) using Seahorse Wave Desktop v2.6.0. Glucose (Sigma #G7021) was added at a final concentration of 10 mM followed by the addition of fumarate (Sigma, F1506) to a final concentration of 10 mM.

### ^13^C_4_-fumaric acid labeling and stable isotope tracing

WT LAC and the Δ*fumC* mutant were grown overnight in LB, then inoculated (1/100) into fresh LB supplemented with ^13^C_4_-fumaric acid (Sigma, 606014, pH 7.0), grown at 37°C to late exponential phase and standardized to an OD_600nm_ of 14. For metabolite extraction, each culture was diluted with 3 volumes of PBS and centrifuged at 2000 x g for 10 minutes at 1°C. The supernatant was discarded, and the pellet was washed with PBS. The pellet (30 μl in volume) was resuspended in a 3:1 methanol:water extraction solution and lysed with 10 freeze-thaw cycles by alternating emersion in liquid nitrogen and a dry-ice/ethanol bath. The debris was removed by centrifugation at 14,000 x g for 5 min at 1°C and the supernatant was stored for analysis. Targeted LC/MS analysis was performed on a Q Exactive Orbitrap mass spectrometer (Thermo Fisher Scientific) coupled to a Vanquish UPLC system (Thermo Fisher Scientific). The Q Exactive operated in polarity-switching mode. A Sequant ZIC-HILIC column (2.1 mm i.d. × 150 mm, Merck) was used for separation of metabolites. Flow rate was set at 150 μl/min. Buffers consisted of 100% acetonitrile for mobile A, and 0.1% NH_4_OH/20 mM CH_3_COONH_4_ in water for mobile B. Gradient ran from 85 to 30% A in 20 min followed by a wash with 30% A and re-equilibration at 85% A. Metabolites were identified based on exact mass within 5 ppm and standard retention times. Relative quantitation was performed based on peak area for each isotopologue. All data analysis was done using MAVEN 2011.6.17. The data in Fig 4 are presented without correction for natural isotope abundance.

### Mouse experiments

All animal experiments were performed following institutional guidelines at Columbia University and Rutgers New Jersey Medical School (NJMS). Animals were housed and maintained at Columbia University Irving Medical Center (CUIMC) and Rutgers Medical Science Building (MSB) under regular rodent light/dark cycles at 18–23°C and fed with irradiated regular chow diet (Purina Cat#5053, distributed by Fisher). Animal health was routinely checked by an institutional veterinary. The animal work protocols (AC-AABD5602 and PROTO202400003) were approved by the Institutional Animal Care and Use Committee (IACUC). Animal experiments were carried out in strict accordance with the recommendations in the Guide for the Care and Use of Laboratory Animals of the NIH, the Animal Welfare Act, and US federal law. C57BL/6 mice, purchased from the Jackson Laboratory (stock number 000664), were acclimated in our facilities. *Ptenl*^*-/-*^ mice (29) were bred in our facilities. The animals were infected at 8 to 10 weeks of age. *In vivo* experiments were performed using roughly 50% female and 50% male animals. During studies, animals were randomly assigned to cages.

### Isolation of murine BMDMs and infection

Murine BMDMs were differentiated as described previously (9). The femurs and tibias were surgically removed from mice. The bone exterior was sterilized with 70% ethanol and the bone marrow was recovered by flushing with phosphate-buffered saline (PBS, Corning, 20-031). The cell suspension was centrifuged for 6 min at 500 x g and resuspended in ACK lysis buffer (Thermo Fisher Scientific, A1049201) to remove the red blood cells. The lysis solution was quenched with PBS, and the cells were centrifuged again and resuspended in DMEM medium (Corning, 10-013-CV) containing 10% heat-inactivated fetal bovine serum (FBS v/v, Sigma, F4135) and 1% penicillin/streptomycin (P/S, v/v, Corning, 30-002-CI)) supplemented with 20 ng/ml rM-CSF (PeproTech #315-02). The rM-CSF-supplemented media was replenished 3 days after isolation, and the BMDMs were mechanically detached on day 7, centrifuged, resuspended in DMEM medium containing 10% FBS (Sigma), and counted using trypan blue stain (Invitrogen, T10282). The cells were seeded at 1 × 10^6^ cells/ml in a 24 well plate (Agilent, 102340-100) and incubated at 37°C with 5% CO_2_ overnight. The cells were infected at a multiplicity of infection (MOI) of 10 for 30 mins and 10 μg ml^−1^ lysostaphin (Sigma) was then added until the desired time point. The cells were washed, detached using TrypLE Express (Life Technologies), serially diluted, and plated on LB agar. Cell viability was determined by counting live cells using trypan blue (Life Technologies).

### Mouse pneumonia model

Mice were infected intranasally with 1 × 10^7^ CFUs of *S. aureus* in 50 μl PBS. PBS alone was used as a control. The mice were sacrificed at 18 h post infection and the BALF and lung were collected. The lung was homogenized through 40 μm cell strainers (Falcon, 352340). Aliquots of the BALF and lung homogenates were serially diluted and plated on LB agar plates and BBL CHROMagar Staph aureus plates (BD) to determine the bacterial burden. The BALF and lung homogenates were spun down and the BALF supernatant was collected for cytokine and untargeted metabolomic analysis. After hypotonic lysis of the red blood cells, the remaining BALF and lung cells were prepared for fluorescence-activated cell sorting (FACS) analysis as described below.

### Flow cytometry of mouse BALF and lung cells

For the identification of immune cell populations, mouse BALF and lung cells were stained in the presence of counting beads (15.45 *μ*m DragonGreen, Bangs Laboratories Inc., FS07F) with LIVE/DEAD stain (Invitrogen, L23105A) and an antibody mixture of anti-CD45-AF700 (BioLegend, 103127), anti-CD11b-AF594 (BioLegend, 101254), anti-CD11c-Bv605 (BioLegend, 117334), anti-SiglecF-AF647 (BD, 562680), anti-MHCII-APC Cy7 (BioLegend, 107628), anti-Ly6C-Bv421 (BioLegend, 128032), and anti-Ly6G-PerCP-Cy5.5 (BioLegend, 127616), each at a concentration of 1:200 in PBS, for 1 h at 4°C. After washing, the cells were stored in 2% paraformaldehyde (PFA, Electron Microscopy Sciences, 15714-S) until analysis on the BD LSRII (BD Biosciences) using FACSDiva v9. Flow cytometry was analyzed with FlowJo v10. Mouse BAL and lung cells were identified as follows:

alveolar macrophages: CD45^+^CD11b^+/−^SiglecF^+^CD11c^+^;

monocytes: CD45^+^CD11b^+^SiglecF^−^MHCII^−^CD11c^−^Ly6G^−^Ly6C^+/−^;

neutrophils: CD45^+^CD11b^+^SiglecF^−^ MHCII^−^CD11c^−^Ly6G^+^Ly6C^+/−^.

### Cytokine analysis

Cytokine concentrations in mouse BALF supernatants were quantified by Eve Technologies (Calgary, Canada) using a bead-based multiplexing technology.

### Semi-targeted metabolomics

Metabolites in the BALF were identified and quantified by high-resolution liquid chromatography mass spectrometry (LC-MS) at the Calgary Metabolomics Research Facility (Calgary, Canada). The metabolites were extracted in a 50% methanol (Alpha Aesar #22909):water (v/v) solution. Sample runs were performed on a Q Exactive HF Hybrid Quadrupole-Orbitrap mass spectrometer (Thermo Fisher Scientific) coupled to a Vanquish UHPLC System (Thermo Fisher Scientific). Chromatographic separation was achieved on a Syncronis HILIC UHPLC column (2.1 mm x 100 mm x 1.7 *μ*m, Thermo Fisher Scientific) using a binary solvent system at a flow rate of 600 μl/min. Solvent A consists of 20 mM ammonium formate pH 3 in mass spectrometry grade water and solvent B consists of mass spectrometry grade acetonitrile with 0.1% formic acid (v/v). The following gradient was used: 0-2 min, 100% B; 2-7 min, 100%–80% B; 7-10 min, 80%–5% B; 10-12 min, 5% B; 12-13 min, 5%–100% B; 13-15 min, 100% B. A sample injection volume of 2 μl was used. The mass spectrometer was run in negative full scan mode at a resolution of 240,000 scanning from 50-750 m/z. Metabolites were identified by matching observed m/z signals (+/−10ppm) and chromatographic retention times to those observed from commercial metabolite standards (Sigma-Aldrich). The data were analyzed using E-Maven and are listed in Tables S8-S9.

### Isolation of RNA from mouse lungs

Total RNA was isolated from excised mouse lungs using TRIzol reagent (Thermo Fisher Scientific, 15596026) according to the manufacturer’s instructions. Residual DNA was then degraded using the TURBO DNA-free Kit (Thermo Fisher Scientific, AM1907) prior to reverse transcription, which was performed using the High-Capacity cDNA Reverse Transcription Kit (Thermo Fisher Scientific, 4374966). Real-time qRT-PCR was performed on the cDNA generated in the previous step using the primers listed in Table S7. The data were analyzed using the ΔΔC_T_ method.

### Human subjects

All human samples (healthy individuals and CF patients) were obtained from adults (22-44 years of age) in the CF program at Yale University under the Yale IRB Protocol 0102012268 and Columbia Protocols AAAR1395. Males and females are represented in an approximate 50–50% ratio. Sputum samples from healthy adult individuals were collected after nebulization with 3% hypertonic saline for five minutes on three cycles. To reduce squamous cell contamination, subjects were asked to rinse their mouth with water and clear their throat. CF subjects expectorated sputum spontaneously for our studies. Expectorated sputum samples were collected into specimen cups and placed on ice. Of the 7 CF patients that were chronically infected with *Staphylococcus aureus* (either MSSA or MRSA), 5 patients were co-infected with other agents, such as *Pseudomonas* (3 patients), *Achromobacter* (1 patient) or human influenza (1 patient). No CF patients were infected with *Aspergillus*, a known itaconate producer. None of the CF patients studied were under CFTR modulator therapy (e.g. Kaleydeco, Orkambi). An informed consent was signed by all subjects.

### Quantification and statistical analysis

We modeled the number of independent experiments required to reach significance between groups using the computer program JMP. This simulation was based on experimental design, preliminary data and past experience. Analyses were performed assuming a 20% standard deviation and equivalent variances within groups. Significance < 0.05 with power 0.8 was used. Experiments in this study were not performed in a blinded fashion. All analyses and graphs were performed using the GraphPad Prism 7a and 8 software. Values in graphs are shown as average ± SEM and data were assumed to follow a normal distribution. For comparison between average values for more than two groups, we performed one-way ANOVA with a multiple posteriori comparison. When studying two or more groups along time, data was analyzed using two-way ANOVA with a multiple posteriori comparison. Differences between two groups were analyzed using a student’s t test for normally distributed data or Mann-Whitney or Kruskal-Wallis test otherwise. Differences were considered significant when a P value under 0.05 (p < 0.05) was obtained. Statistical details of experiments are indicated in each figure legend. No data points were excluded.

## Data availability

All data discussed in this study are presented in the published article and its supplementary files. RNA-Seq data are deposited under accession number GSE276380.

## Acknowledgments

We thank Drs. Xiaoyuan Yang and Guoan Zhang from the Proteomics and Metabolomics Core Facility at Weill Cornell Medicine for the staple isotope tracing analysis. We also thank Dr. Min Li (Shanghai Jiao Tong University School of Medicine, China) for sharing *S. aureus* strain FPR3757, Dr. Clemente J. Britto (Yale University) for providing the sputum samples, which were initially used in our previous study (9), and Drs. Paul Fey and Itidal Reslane (University of Nebraska) for insightful discussions. This study is supported by NIH grant K99/R00 HL157550 to T.W.F.L., NIH grant R35GM146776 to S.A.R., NIH grant R01 HL170129 to A.P., and NIAID 1R01AI139100-01, NSF 1750624 and USDA MRF project NE−1028 awards to JMB. This study is also supported by NIH grant S10RR027050 to the Columbia Center for Translational Immunology (CCTI) Flow Cytometry Core. B.P.H., T.P.S. and A.H. are supported by NHMRC Investigator grants (Australia) APP1196103, GNT1194325, and APP2018880. C.W. is supported by the National Key R&D Program of China (2022YFA1304700). I.A.L. is supported by an Alberta Innovates Translational Health Chair. The Calgary Metabolomics Research Facility (CMRF) is supported by the International Microbiome Centre and the Canada Foundation for Innovation (CFI-JELF 34986).

## Author contributions

Y-T.C., Z.L.^2,#^ and D.F. contributed equally, performing experiments, analyzing data, and editing the manuscript. Z.L.^4^ synthesized the fumarate analogue probe. Z.L.^2,#^ and C.W. specifically performed and analyzed the chemoproteomic experiments with the fumarate analogue probe. S.G.G. and A.H. performed *in silico* analyses on the conservation of *fumC* in *S. aureus* isolates. H.H. performed the homology modeling. C.B. and P.P. provided whole genome sequences of a collection of *S. aureus* isolates from cystic fibrosis patients. G.R. and J.M.B. measured *S. aureus* aconitase activity. N.P. constructed the Δ*sucA* and Δ*sucD* mutants in the USA300 LAC background and R.W. performed *in vivo* experiments and analyzed data. I.R.M. constructed the bacterial Δ*fumC* mutant and complemented strain. G.L. and A.Z.B. isolated murine bone marrow-derived macrophages. S.S.S. and S.H.S. analyzed the bulk RNA-seq data. M.D. performed the LC-MS and S.A.R. provided the data on the relative quantification of fumarate and itaconate in the sputum of patients with cystic fibrosis versus healthy controls. F.A.B.O. performed confocal imaging on biofilms. I.A.L., R.S., T.P.S., B.P.H., J.M.B., A.H., I.R.M., A.T., S.A.R. and C.W. contributed to the experimental design and discussions. A.P. co-wrote and edited the manuscript. T.W.F.L. proposed the central hypothesis, performed experiments, analyzed/interpreted the data, and wrote/edited the manuscript.

## Inclusion and ethics statement

All collaborators, local and international, have made significant contributions to the study design, data collection, and analysis, and are listed as co-authors on this manuscript. This research was not restricted in the setting of the researchers and does not result in stigmatization or discrimination. Local and regional research relevant to our study was considered and included in citations.

## Figure legends

**Extended Data Figure 1.**
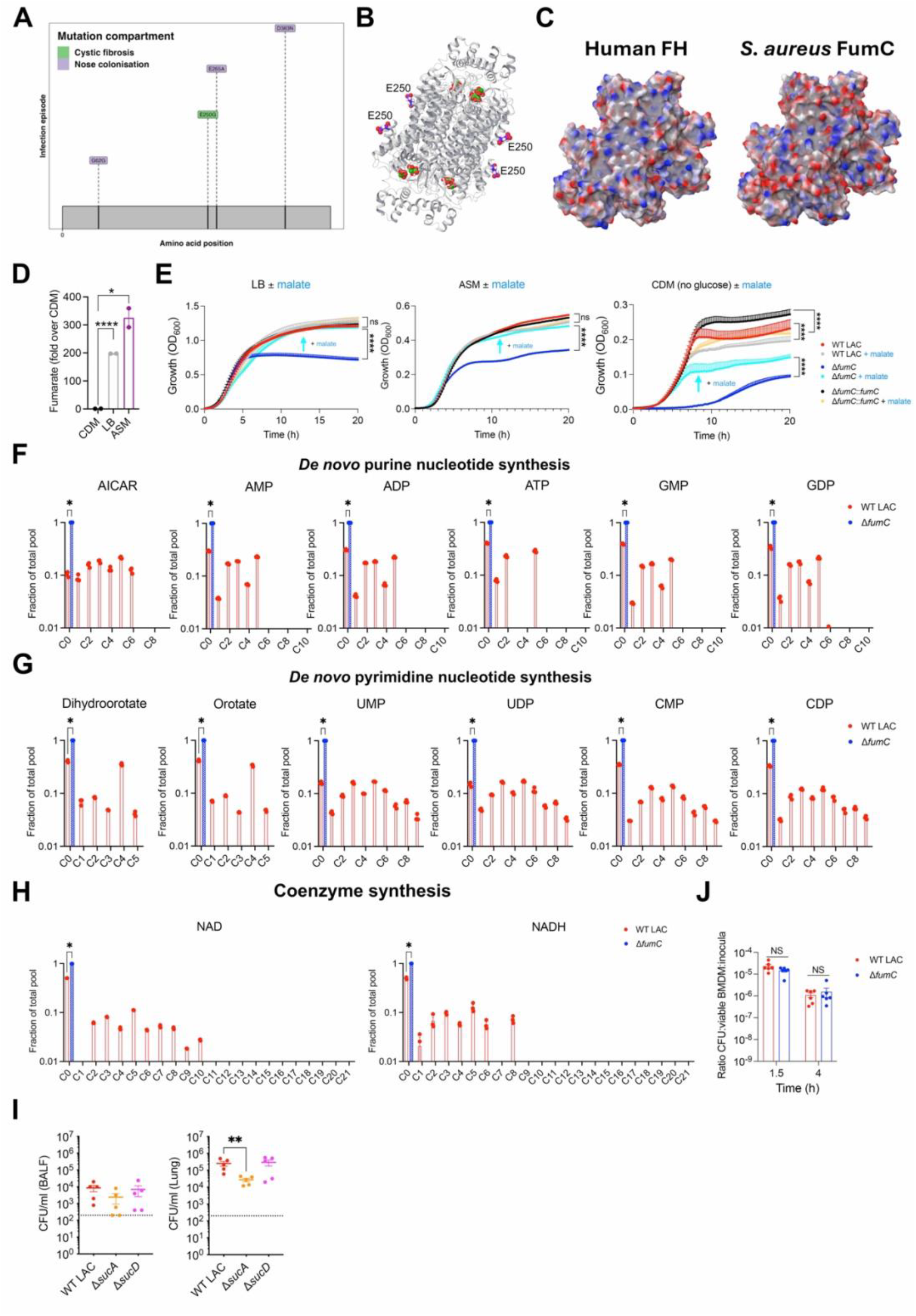
FumC is essential for bacterial metabolic fitness. (**A**) Map of *fumC* showing the position of *de novo* mutations in both pulmonary isolates and nasal colonizers. (**B**) Simulated structure of *S. aureus* FumC tetramer depicting the position of the SNP E250G (carbon atoms in purple) with respect to the catalytic site. Citrate and malate ligands bound to the active and B sites respectively, are shown with carbon atoms in green. Ligand positions were created by alignment with PDB ID: 6OS7, and 1FUR. (**C**) Protein surface colored by electrostatic potential, where red symbolizes negative charge, and blue positive charge. The left panel displays human fumarate hydratase (PDB: 5UPP). The right panel illustrates the constructed model of *S. aureus* FumC. **(D)** Relative level of fumarate in chemically defined media (CDM), Luria-Bertani broth (LB) and artificial sputum media (ASM), n = 2 per condition. **(E)** Growth curve of WT LAC, the Δ*fumC* mutant and complemented strain in LB (left panel), ASM (middle panel) or CDM (right panel) supplemented with or without 10 mM malate; n = 3 biological samples in triplicate (3 independent experiments). **(F-H)** ^13^C-fumarate labeling of metabolites involved in the **(F)** *de novo* purine nucleotide synthesis, **(G)** *de novo* pyrimidine nucleotide synthesis, or **(H)** coenzyme synthesis in WT LAC and the Δ*fumC* mutant; n = 3 biological replicates (1 independent experiment). **(I)** Bacterial burden from the BALF (left panel) and lung (right panel) of BL/6 infected with WT LAC and the Δ*sucA* and Δ*sucD* mutants (n = 5 per condition). The dotted line indicates the limit of detection. **(J)** Intracellular bacterial survival at 1.5 h and 4 h post-infection in BMDMs from WT BL/6 mice relative to the number of viable BMDMs and bacterial inocula; n = 2 independent experiments. Data are shown as mean ± SEM. Statistical significance is determined by two-tailed t-student test with FDR correction for (E-I) and Mann-Whitney non-parametric U test for (D), *p < 0.05, **p < 0.01, ***p < 0.001 and ****p < 0.0001.

**Extended Data Figure 2.**
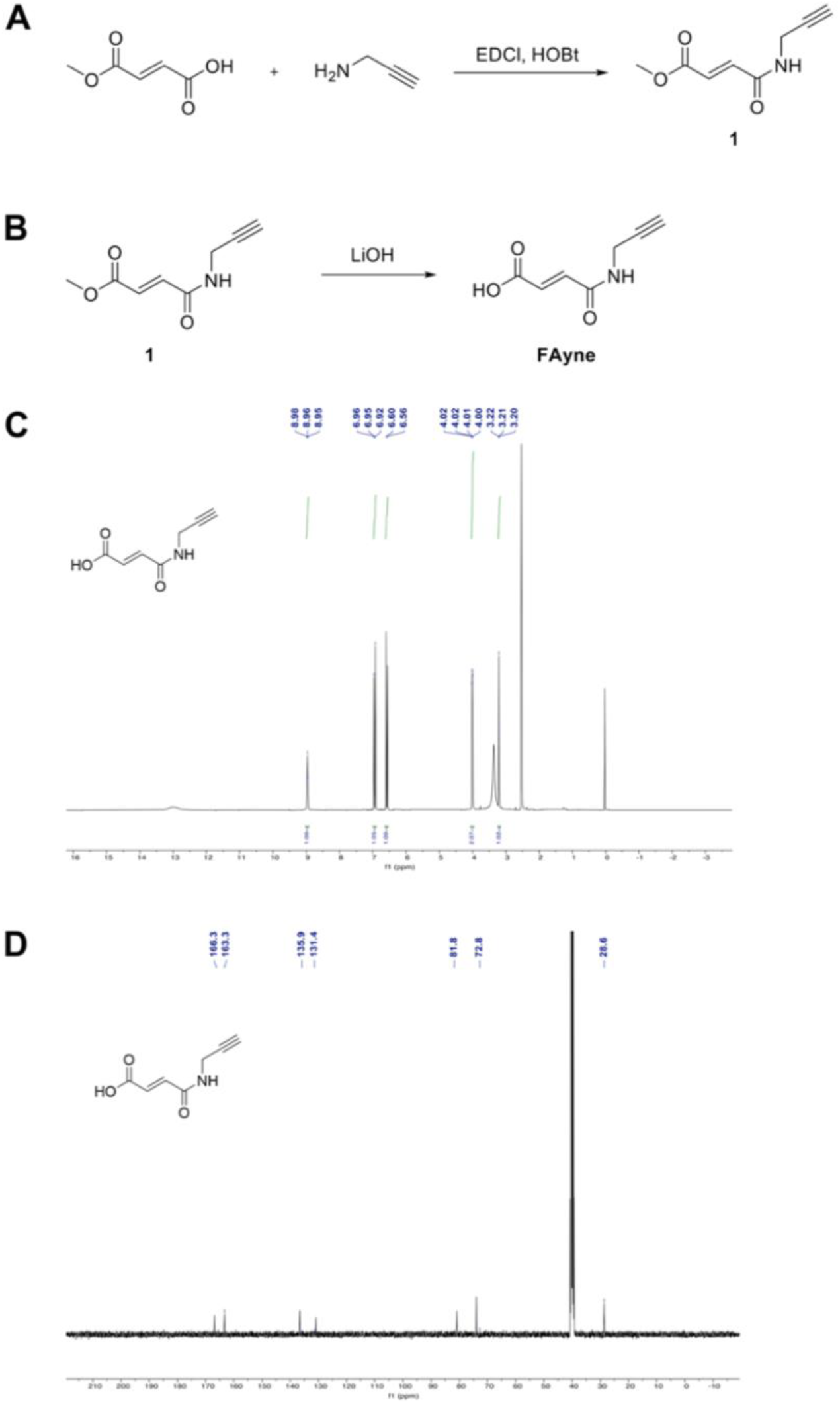
Synthesis of the FA-alkyne probe. **(A-B)** Chemical reactions for the production of **(A)** compound 1 and **(B)** the FA-alkyne probe. **(C-D)** Validation of the structure of the FA-alkyne probe by **(C)** proton nuclear magnetic resonance (^1^H NMR) and **(D)** carbon-13 nuclear magnetic resonance (^13^C NMR).

